## Supplemental Information for "Asymmetric Structure of the Native *Rhodobacter sphaeroides* Dimeric LH1-RC Complex"

K. Tani, R. Kanno et al.

**Supplementary Table 1 Cryo-EM data collection, refinement and validation statistics.**

| | Native dimeric LH1-RC complex (EMDB-32192, PDB ID: 7VY2) | Monomeric LH1-RC ΔU complex (EMDB-32193, PDB ID: 7VY3) |
|---|---|---|
| **Data collection and processing** | | |
| Magnification | 96000 | 96000 |
| Voltage (kV) | 300 | 300 |
| Electron exposure (e–/Å$^2$) | 40 | 40 |
| Defocus range (μm) | –1.0 to –2.9 | –0.8 to –2.7 |
| Pixel size (Å) | 0.820 | 0.820 |
| Symmetry imposed | C1 | C1 |
| Initial particle images (no.) | 164496 | 337006 |
| Final particle images (no.) | 124916 | 124589 |
| Map resolution (Å) | 2.8 | 2.6 |
| FSC threshold | 0.143 | 0.143 |
| Map resolution range (Å) | 369–2.8 | 295–2.6 |
| **Refinement** | | |
| Initial model used (PDB code) | 7F0L | 7F0L |
| Model resolution (Å) | 3.1 | 2.6 |
| FSC threshold | 0.5 | 0.5 |
| Model resolution range (Å) | 220–2.8 | 125–2.6 |
| Map sharpening *B* factor (Å$^2$) | –61 | –70 |
| Model composition | | |
| Non-hydrogen atoms | 46314 | 18831 |
| Protein residues | 4534 | 1846 |
| Ligands | 201 | 86 |
| *B* factors (Å$^2$) | | |
| Protein | 46.1 | 29.7 |
| Ligand | 52.7 | 31.0 |
| R.m.s. deviations | | |
| Bond lengths (Å) | 0.005 | 0.006 |
| Bond angles (°) | 2.667 | 2.676 |
| Validation | | |
| MolProbity score | 1.83 | 1.83 |
| Clashscore | 13.13 | 10.91 |
| Poor rotamers (%) | 1.62 | 2.30 |
| Ramachandran plot | | |
| Favored (%) | 97.81 | 97.99 |
| Allowed (%) | 2.19 | 2.01 |
| Disallowed (%) | 0.00 | 0.00 |

**Supplementary Table 2  Comparison of the distances of His–BChl(Mg) and BChl(Mg)–BChl(Mg) in LH1, LH2 and RC special pairs from various phototrophic bacteria.**

| | Distance of His(Nε2) to BChl–Mg (Å) | | Distance of Mg–Mg (Å) | | |
|---|---|---|---|---|---|
| LH1 or LH2 | α | β | Long | Short | PDB ID |
| ***Rba. sphaeroides* (Dimer)** | **2.59** | **2.21** | **9.49** | **8.34** | **7VY2** |
| ***Rba. sphaeroides* (ΔU)** | **2.52** | **2.24** | **9.54** | **8.23** | **7VY3** |
| *Rba. sphaeroides* (Monomer) | 2.58 | 2.20 | 9.55 | 8.37 | 7F0L |
| *Rsp. rubrum* (LH1) | 2.27 | 2.03 | 9.34 | 8.51 | 7EQD |
| *Rps. palustris* (LH1-W) | 2.93 | 2.71 | 9.61 | 8.2 | 6Z5S |
| *Tch. tepidum* (LH1) | 2.19 | 2.19 | 8.88 | 8.72 | 5Y5S |
| *Trv.* strain 970 (LH1) | 2.33 | 2.31 | 8.90 | 8.46 | 7C9R |
| *Blc. viridis* (LH1) | 2.54 | 2.25 | 8.8 | 8.5 | 6ET5 |
| *Rfx. castenholzii* (B880) | 2.32 | 2.29 | 9.5 | 9.3 | 5YQ7 |
| *Rbl. acidophilus* (B850) | 2.34 | 2.34 | 9.5 | 8.8 | 1NKZ |
| *Phs. molischianum* (B850) | 2.27 | 2.32 | 9.2 | 8.9 | 1LGH |

| RC (special pair) | L-subunit | M-subunit | BChl *a* (L)–BChl *a* (M) | |
|---|---|---|---|---|
| ***Rba. sphaeroides* (Dimer:A)** | **2.24** | **2.15** | **7.80** | **7VY2** |
| ***Rba. sphaeroides* (Dimer:B)** | **2.20** | **2.15** | **7.77** | **7VY2** |
| ***Rba. sphaeroides* (ΔU)** | **2.24** | **2.37** | **7.76** | **7VY3** |
| *Rba. sphaeroides* (RC-only) | 2.27 | 2.06 | 7.84 | 2J8C |
| *Rsp. rubrum* | 2.09 | 2.12 | 7.76 | 7EQD |
| *Rps. palustris* | 2.73 | 2.74 | 7.69 | 6Z5S |
| *Tch. tepidum* | 2.17 | 2.19 | 7.87 | 5Y5S |
| *Trv.* strain 970 | 2.33 | 2.31 | 7.65 | 7C9R |
| *Blc. viridis* | 2.36 | 2.35 | 7.83 | 6ET5 |

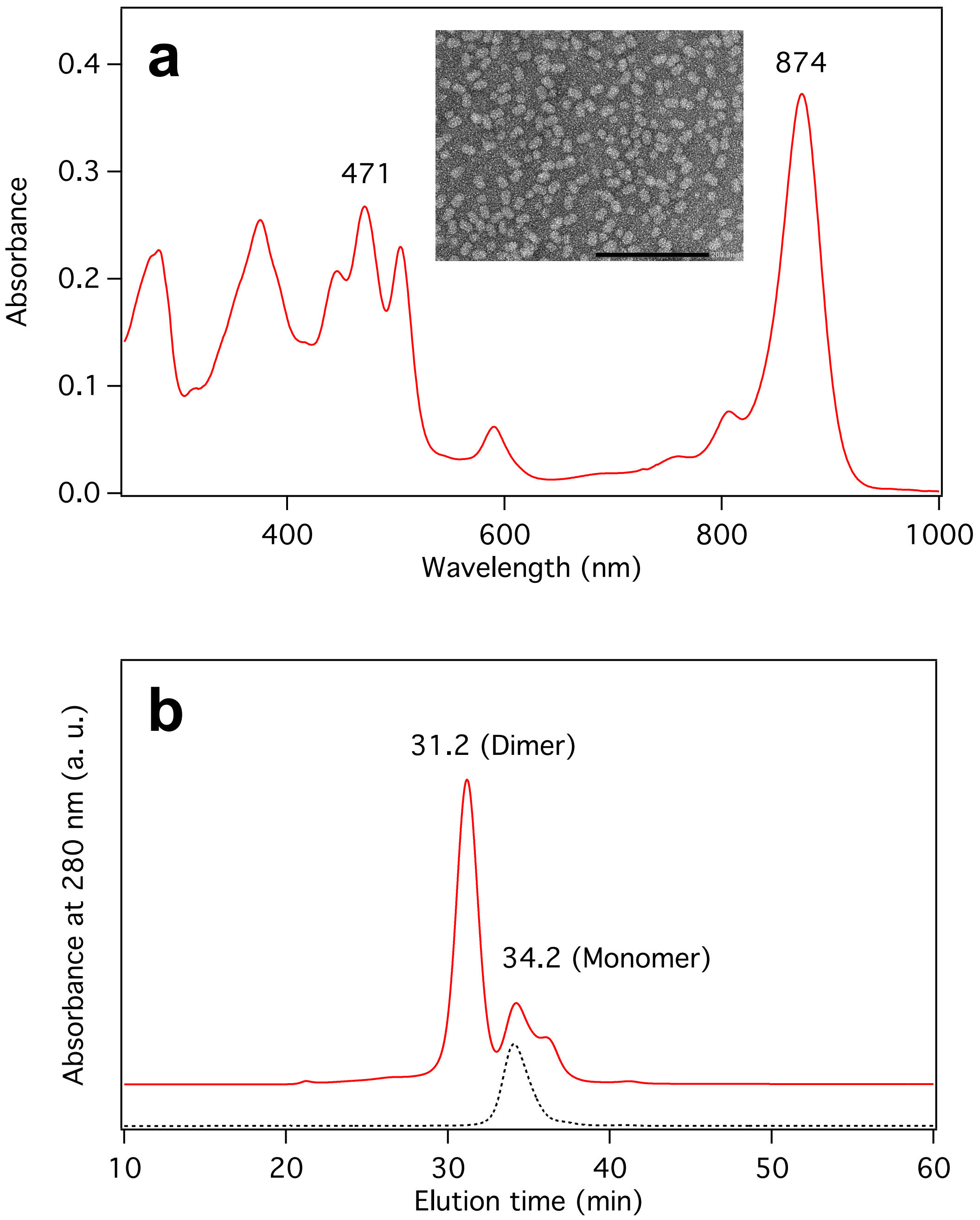

**Supplementary Fig. 1 Absorption spectrum and gel-filtration chromatogram of the native *Rba. sphaeroides* IL106 dimeric LH1-RC complex.** **(a)** Absorption spectrum of the purified dimeric LH1-RC at room temperature. Inset shows negatively stained LH1-RC particles obtained with 0.04 mg/mL LH1-RC in 20mM Tris-HCl (pH7.5) containing 0.05% DDM. Scale bar: 200 nm. **(b)** Gel-filtration chromatogram of the dimer-rich LH1-RC purified from DEAE chromatography using TSKgel G4000SW (7.5 mm I.D. × 60 cm, TOSOH) with 20 mM Tris-HCl buffer (pH 7.5) containing 200 mM NaCl and 0.05 % w/v DDM at flow rate of 1 mL/min at room temperature. Dashed black line shows the chromatogram of a highly purified monomeric LH1-RC.

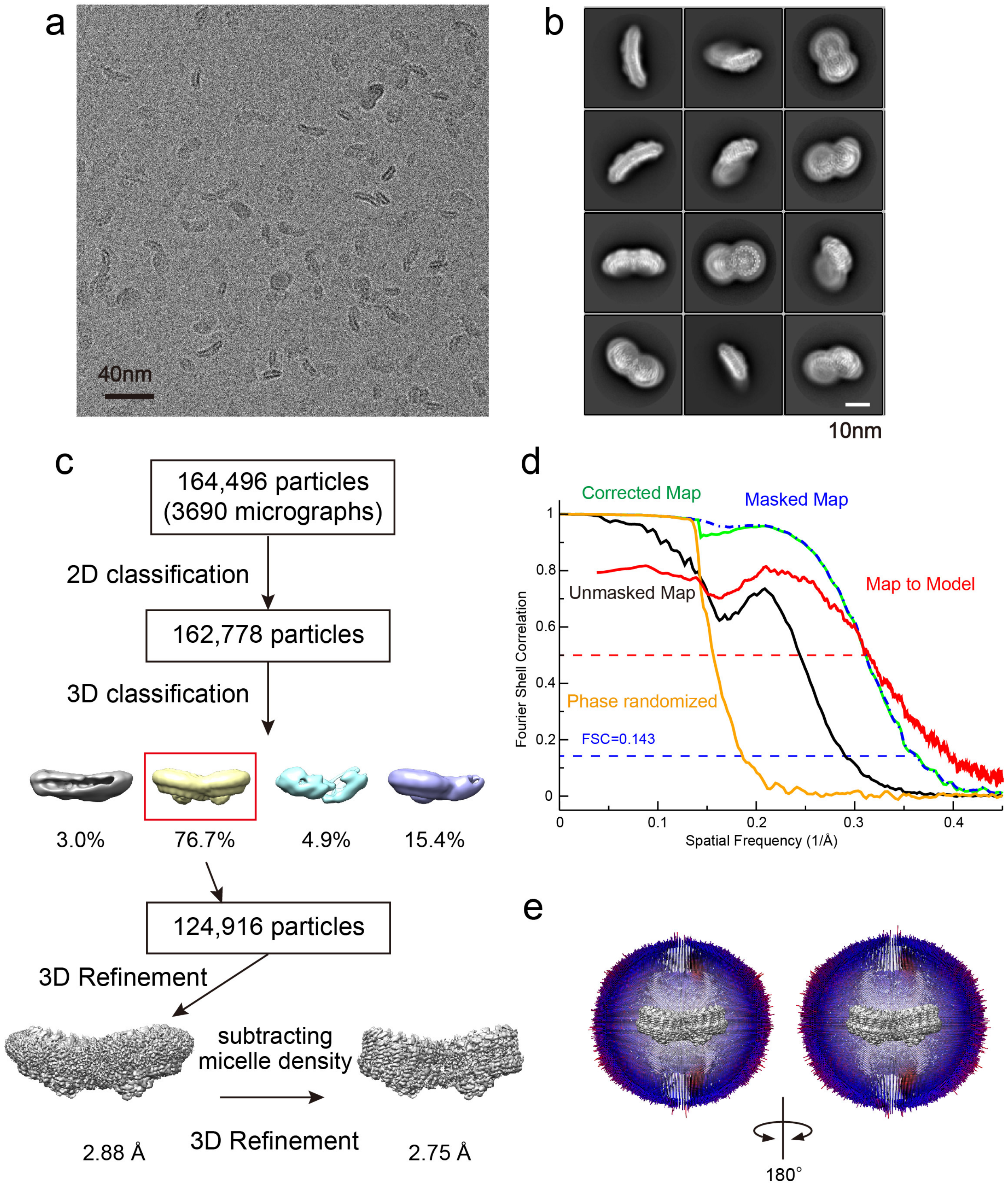

**Supplementary Fig. 2 Structure determination of the native *Rba. sphaeroides* IL106 dimeric LH1-RC complex. (a)** A representative cryo-EM micrograph. **(b)** Representative 2D class averages processed from the micrographs of LH1-RC. **(c)** Image processing flow of 3D classification and reconstruction. **(d)** The Fourier shell correlation (FSC) plot of the cryo-EM map (unmasked: black, masked: blue, phase randomized corrected: green, phase randomized: orange) and the FSC plot of the model versus the final map (red) are superimposed. **(e)** Angular distribution of reconstructed particles in the C1 map of dimeric LH1-RC complex. For clarity, the front half of angular distribution is removed.

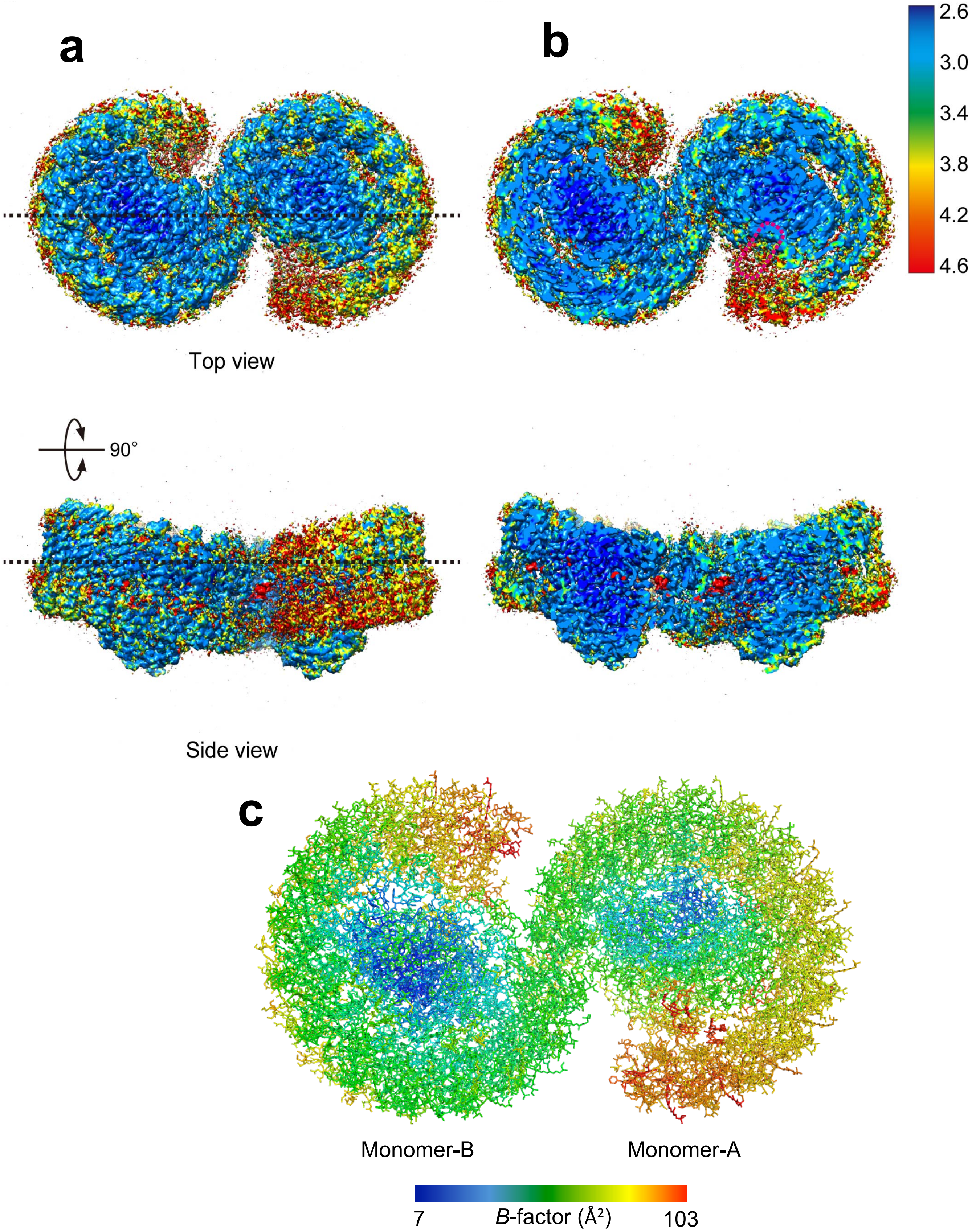

**Supplementary Fig. 3  Local resolution and *B*-factor distributions of the native dimeric LH1-RC structure.** **(a)** Top (periplasm) and side views of local resolution distributions. Dotted lines indicate cross sections. **(b)** Central crossing section views of local resolution distributions for the panels on the left-hand sides. The map color codes represent the resolutions from 4.6 Å (red) to 2.6 Å (blue) as indicated in the color bars. **(c)** Top view of *B*-factor distribution from periplasmic side. The color codes represent the *B*-values indicated in the color bar.

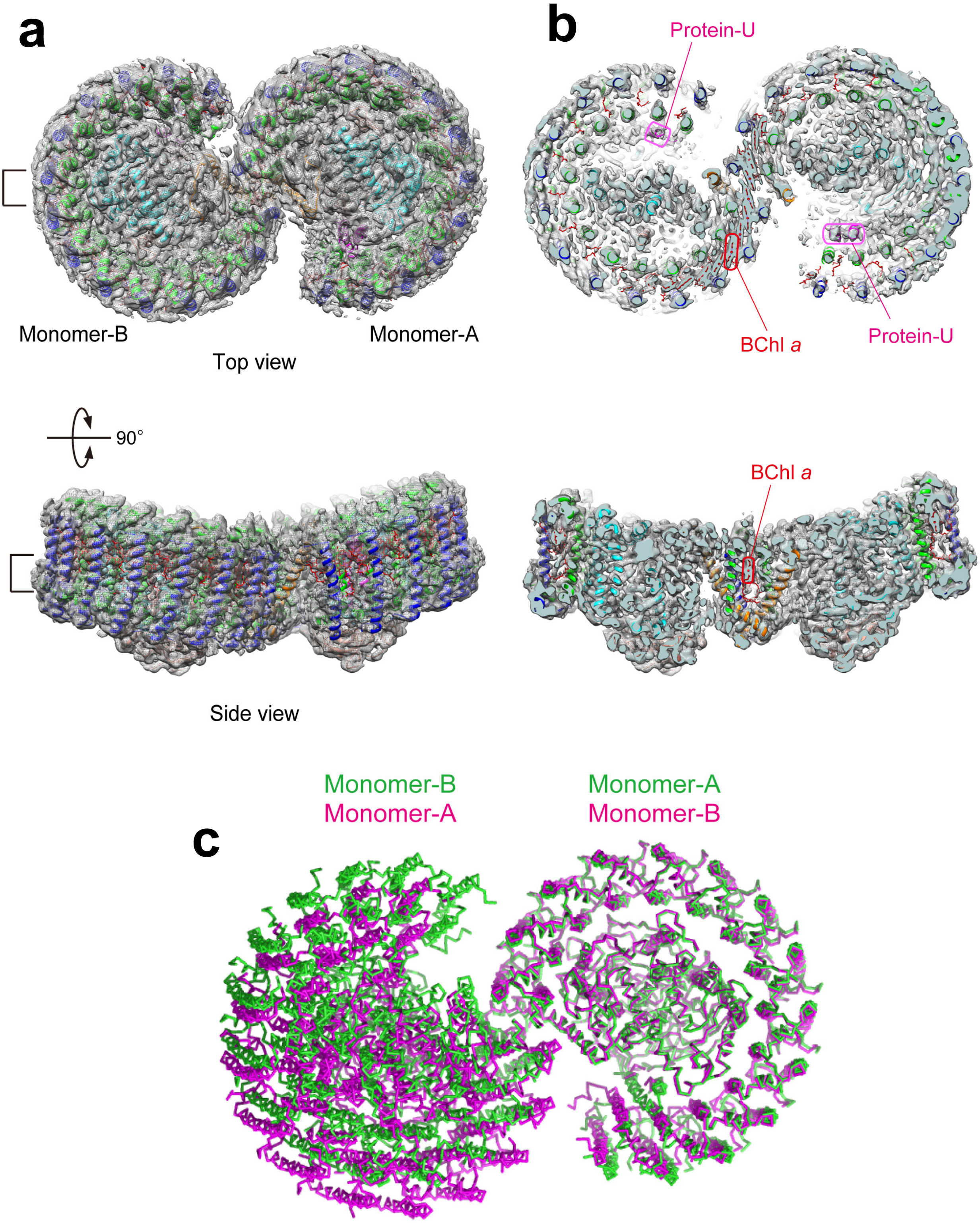

**Supplementary Fig. 4 Overall structure of the native dimeric LH1-RC complex.** Cartoon representation of the complex with cryo-EM density map shown in gray mesh. **(a)** Top view from periplasmic side and side view parallel to the membrane plane. Each bracket indicates the slab region. **(b)** Top and side views of the slabs in (a) along and perpendicular to the membrane plane. The color codes corresponding to polypeptides and BChl *a* are the same as in Fig.1. Regions indicated by pink circles correspond to BChl *a*. **(c)** Overlap view of the two dimers by superposition of Cα carbons of the RC L-subunit in monomer-A (green) with those in monomer-B (180° rotated, magenta), showing asymmetric positions between the two monomers.

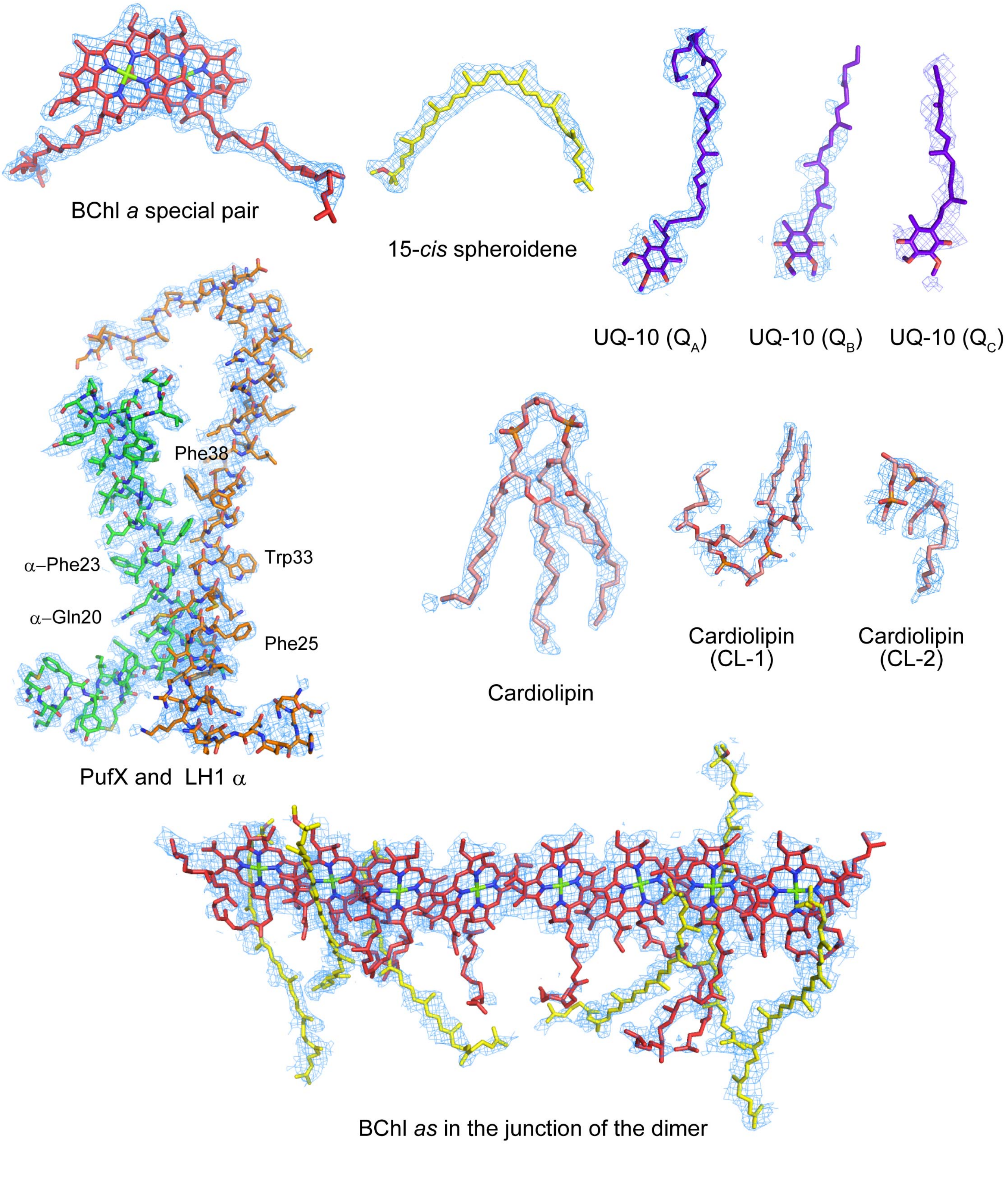

**Supplementary Fig. 5 Cryo-EM densities and structural models in the dimer of the *Rba. sphaeroides* IL106 LH1-RC complex.** The color codes of polypeptides are the same as in Fig. 1. The density maps are shown at a contour level of 4.0σ.

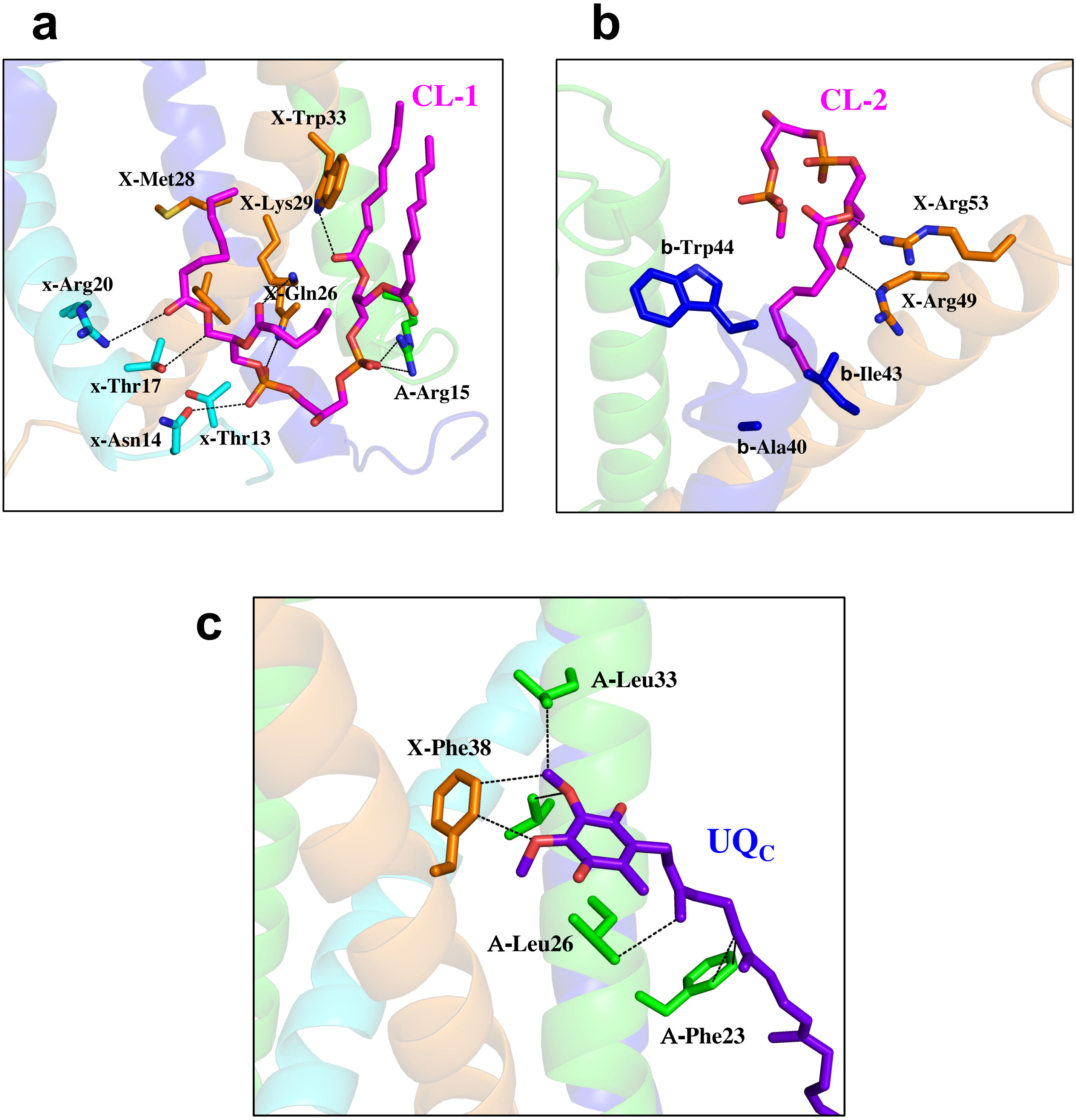

**Supplementary Fig. 6 Interactions of the cardiolipin (CL) and UQ molecules in the dimer junction with nearby polypeptides. (a)** Close contacts (<4.0 Å) between CL-1 (magenta sticks) and two PufX polypeptides (orange and cyan with chains ID: X and x) on the cytoplasmic side. Dashed lines indicate possible hydrogen bondings. **(b)** Close contacts (<4.0 Å) of CL-2 (magenta sticks) with a PufX (orange, chain ID: X) and a β-polypeptide (blue, chain ID: b) belonging to another monomer on the periplasmic side. **(c)** Close contacts (<4.0 Å, dashed lines) of $UQ_C$ (purple sticks) with a PufX (orange, chain ID: X) and an α-polypeptide (green, chain ID: A) in the transmembrane region.

**a**

```
               31 34
PufX:     QMMKGAGWAGGVFFGT
α-chain:  RRVFVAQGVFLFLLAV
              18 21
```

**b**

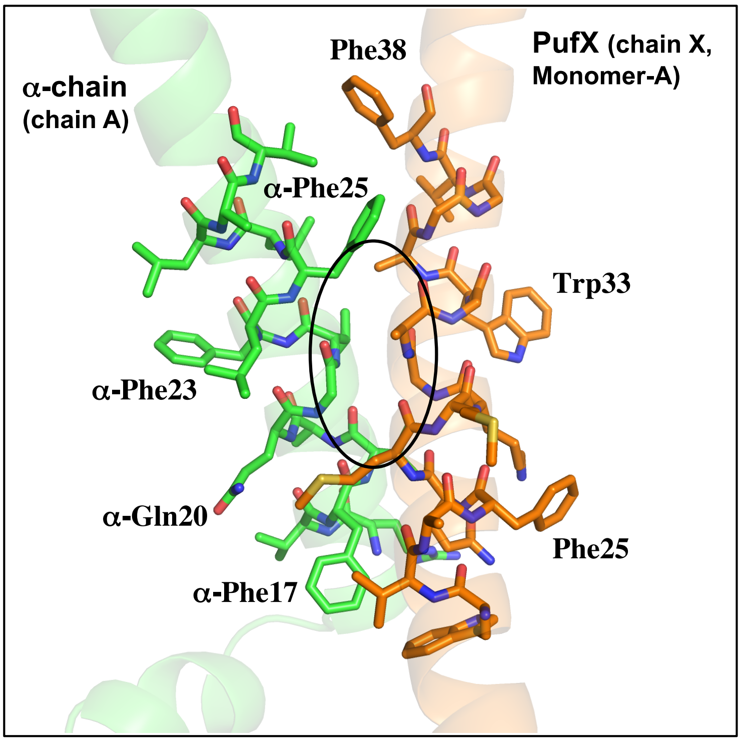

**c**

```
                           10        20        30        40        50        60        70        80
Rba. sphaeroides   MADKTIFNDHLNTNPKTNLRLWVAFQMMKGAGWAGGVFFGTLLLIGFFRVVGRMLPIDENPAPAPNITGALETGIELIKHLV
Rba. azotoformans  MADKTIFDDHLKTNPKTNLRLWVAFQMMKGAGWAGAVFFGTLLMIGFFRVLGRALPIDENPAPAPNLTGALETGIELIKHLV
Rba. veldkampii    MAEKHYLDGATKVG----MATMGAAAMGKGMGITAVVFFGTVFFVVALAFIGQFLPDRSREAPYPNTIFQVNDIDGTVDGKYTRFAN
Rba. capsulatus    MSMFDKPFDYENGS-KFEMGIWIGRQMAYGAFLGSIPFLLGLGLVLGSYGLGLMLPERAHQAPSPYTTEVVVQHATEVV
Rba. blasticus     MAEYNYSHEPNAVI---NLRVWALGQMVWGAFLAAVGVVVVICLLVGTYLAGLLLPEQSKQAPSPYGALEIVQTIDVA
                   *                         *  *                     *  **     ** *
```

**d**

```
                            10        20        30        40        50
Rba. sphaeroides    -----MSKFYKIWMIFDPRRVFVAQGVFLFLLAVMIHLILLSTPSYNWLEISAAKYNRV
Rba. azotoformans   -----MAKFYKIWMIFDPRRVFVAQGVFLFLLAVMIHLILLSTPSYNWLEISAAKYNRVAVAE
Rba. veldkampii     -----MSKFYKIWLIFDPRRVFVAQGVFLFLLAVMIHLMLLSNPGFNWLDISGVKYERVAAE
Rba. capsulatus     -----MSKFYKIWLVFDPRRVFVAQGVFLFLLAVLIHLILLSTPAFNWLTVATAKHGYVAAAQ
Rba. blasticus      -----MSKFYKIWQVFDPRRVFVAQGVFLFLLAVMIHLILLSKPDYNWLDVGTAKYGRGEAAAVVTP
Tch. tepidum        MFTMNAN-LYKIWLILDPRRVLVSIVAFQIVLGLLIHMIVLST-DLNWLDDNIPVSYQALGKK
Trv. strain 970 A1  MNAKSFDGMHKLWMIMNPVSTLWAIFIFQIFLGLLIHMVVLSS-DLNWHDDQIPVGYQLQGETLPVNLEMKAAQ
Trv. strain 970 A3  MNAKSFDGMHKLWMIMNPVSTLWAIFIFQIFLGLLIHMVVLSS-DLNWHDDQIPVGYQLQGETLPVNLEMKAALKDAQ
Blc. viridis        MATEYRTASWKLWLILDPRRVLTALFVYLTVIALLIHFGLLSTDRLNWWEFQRGLPKAA
Rsp. rubrum         --------MWRIWQLFDPRQALVGLATFLFVLALLIHFILLSTERFNWLEGASTKPVQTSMVMPSSDLAV
                                *    *                 **   **    **
```

**Supplementary Fig. 7 Interactions between PufX and α-polypeptide in the transmembrane region. (a)** Sequence alignment of the crossing interface between the two polypeptides with a Gly- and Ala-rich stretch (red fonts) in PufX facing to a short segment (VAQGV, red fonts) in the α-polypeptide. **(b)** Contact interface (elliptical box) between the two polypeptide grooves formed by residues with the smallest side chains (essentially Gly) whereas most bulky side chains point outward the interface. **(c)** Sequence alignments of the PufX from five *Rhodobacter* species showing a conserved stretch rich in Gly and Ala (red fonts). **(d)** Sequence alignments of the α-polypeptides for typical purple bacteria showing a conserved VAQGV segment in the *Rhodobacter* species**.** Symbol (*) denotes identical residues.

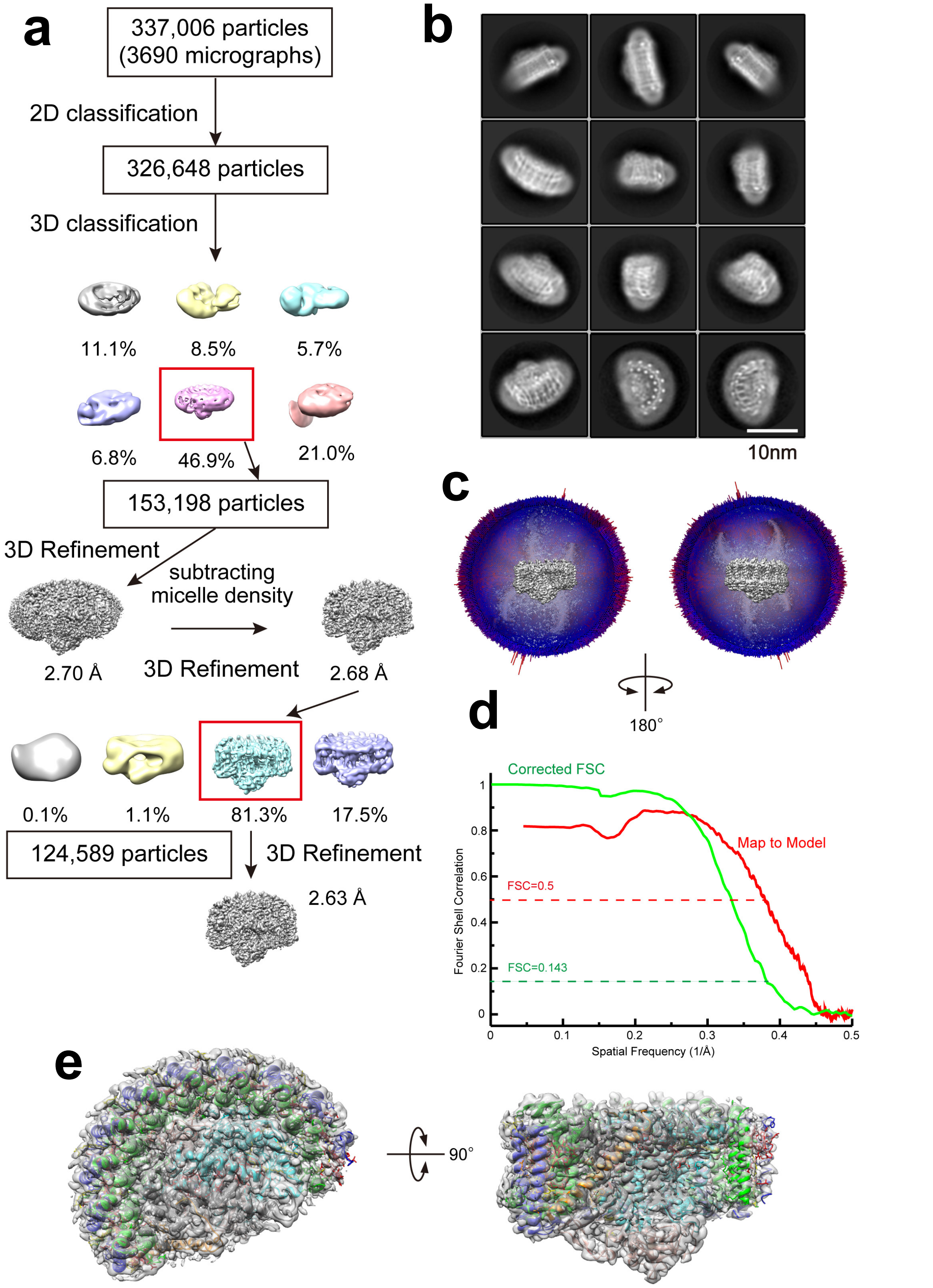

**Supplementary Fig. 8 Structure determination of the monomeric LH1-RC ΔU complex. (a)** Image processing flow of classification and 3D reconstruction. **(b)** Representative 2D class averages of ΔU LH1-RC monomer. **(c)** Angular distribution of reconstructed particles in the C1 map of ΔU LH1-RC complex. For clarity, the front half of angular distribution is removed. **(d)** Fourier shell correlation (FSC) plot of the cryo-EM map (phase randomized corrected: green) and the FSC plot of model versus the final map (red) are superimposed. **(e)** Cartoon representation of the complex with density shown in gray mesh. Top view from periplasmic side and side view parallel to the membrane plane.

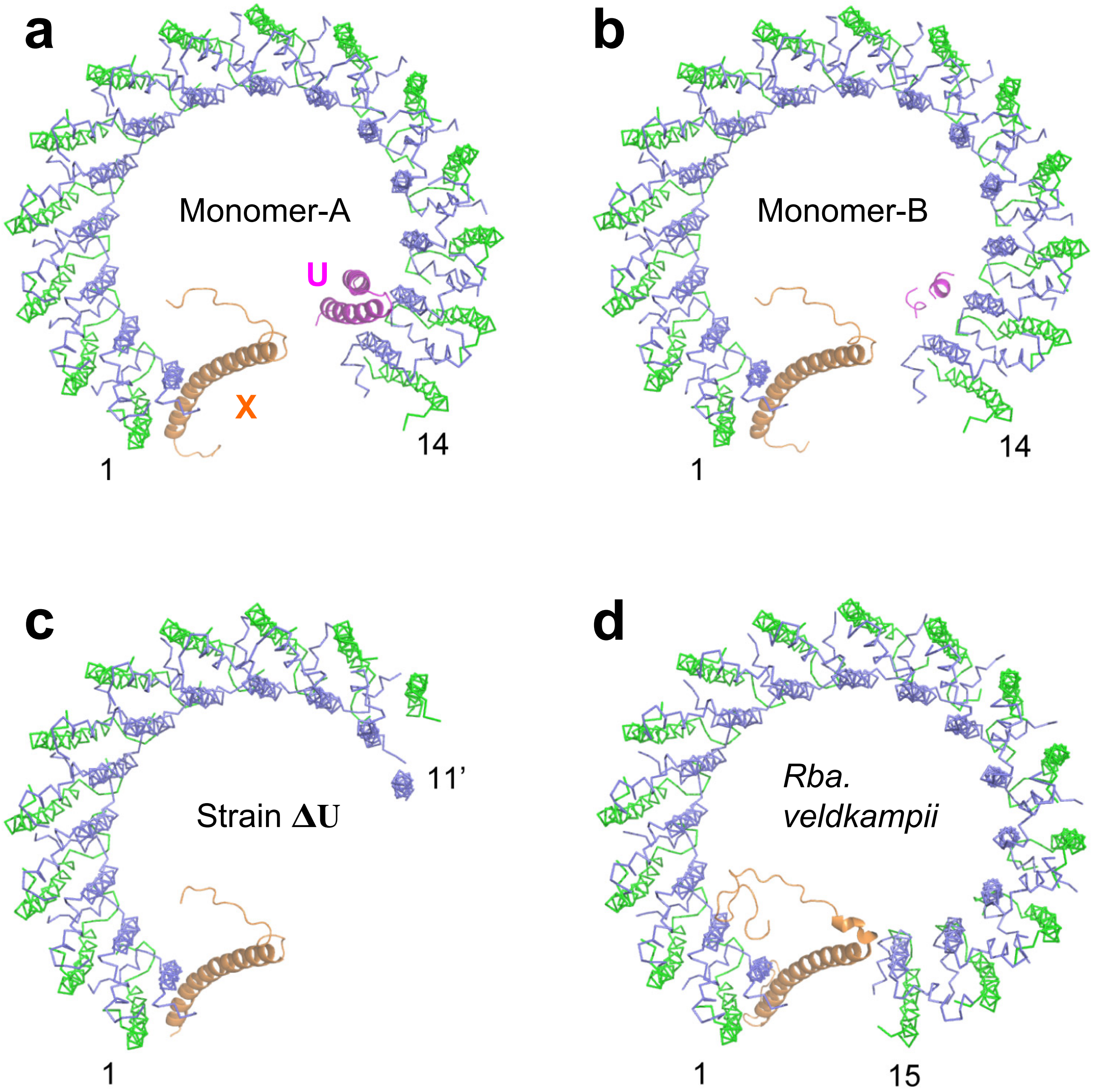

**Supplementary Fig. 9 Comparison of the LH1 ring structure determined in this work with those reported in other LH1-RC-PufX structures.** Top views of superposition of the Cα carbons of the LH1 αβ–polypeptides between the structure in this work (LH1-α, green ribbon; LH1-β, slate-blue ribbon; PufX, orange cartoon; Protein-U, magenta cartoon). **(a)** Monomer-A of the dimer in this work. **(b)** Monomer-B of the dimer in this work. **(c)** Protein-U-lacking monomeric LH1 from strain ΔU (this work). **(d)** Monomeric LH1 of *Rba. veldkampii* (PDB: 7DDQ).

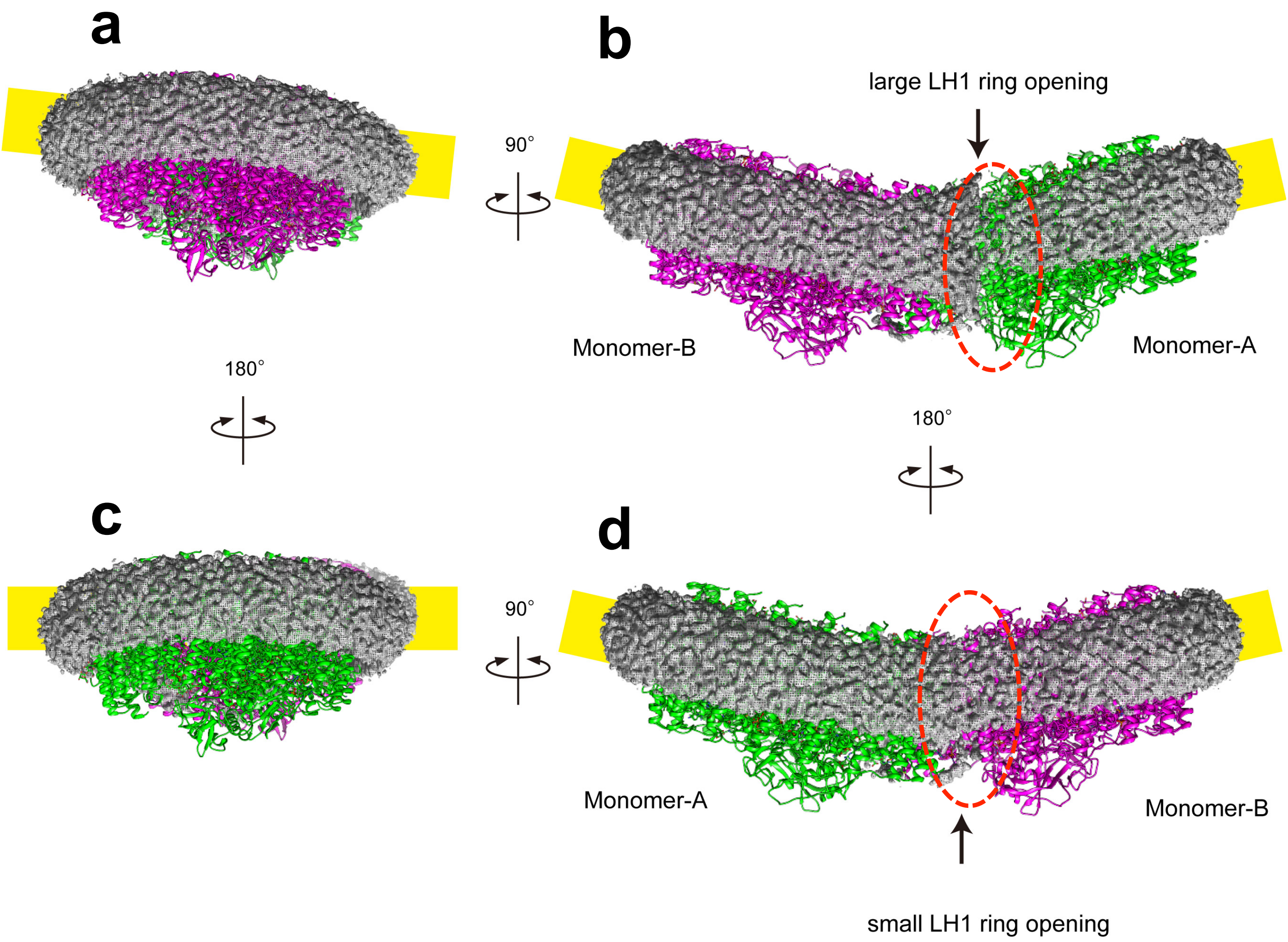

**Supplementary Fig. 10 Asymmetric distributions of lipid/micelles around the LH1-RC dimer.** Monomer-A, green cartoon; monomer-B, magenta cartoon; density maps for lipid/micelles, gray mesh; deduced membrane boundary, yellow box. **(a)** Side view from monomer-B along the long axis of dimer. The deduced membrane boundary is distorted. **(b)** Side view of the dimer with monomer-A on the right-hand side showing a larger LH1-ring opening (red dashed ellipse). The lipid/micelle belt is disconnected and the deduced membrane boundary is kinked by an angle of ~158°. **(c)** Side view from monomer-A along the long axis of dimer. The deduced membrane boundary is nearly flat. **(d)** Side view of the dimer with monomer-B on the right-hand side showing a smaller LH1-ring opening (red dashed ellipse). The lipid/micelle belt is continued and the deduced membrane boundary is kinked by an angle of ~152°.
